# Default mode and motor networks facilitate early learning of implicit motor sequences: a multimodal MR spectroscopy and fMRI study

**DOI:** 10.1101/2024.12.07.627311

**Authors:** Joshua Hendrikse, Emily Brooks, Sarah Wallis, Dylan Curtin, Nigel C. Rogasch, Murat Yücel, Mana Biabani, Charlotte J Stagg, Mark Bellgrove, Richard McIntyre, Chao Suo, James Coxon

**Affiliations:** School of Psychological Sciences & Turner Institute for Brain and Mental Health, Monash University, Melbourne, Victoria, Australia; Hopwood Centre for Neurobiology, Lifelong Health Theme, South Australian Health and Medical Research Institute (SAHMRI); Discipline of Physiology, School of Biomedicine, University of Adelaide; Queensland Institute of Medical Research Berghofer (QIMR), Brisbane; Wellcome Centre for Integrative Neuroimaging, FMRIB, Nuffield Department of Clinical Neurosciences, University of Oxford, UK; Medical Research Council Brain Network Dynamics Unit, Nuffield Department of Clinical Neuroscience, University of Oxford, UK

**Keywords:** Motor learning, cardiovascular exercise, neuroplasticity, micro-consolidation, hippocampus, motor cortex, GABA, MR spectroscopy

## Abstract

Learning new motor skills is a fundamental process that involves the sequencing of actions. Skill develops with practice and time, and manifests as performance that is fast and accurate. Although we know that learning can occur through an *implicit* process in the absence of conscious awareness, and across multiple temporal scales, the precise neural mechanisms mediating implicit motor sequence learning remain poorly understood. Similarly, the capacity for interventions with known influence on learning and memory, such as cardiovascular exercise, to facilitate implicit learning is yet to be clearly established. Here, we investigated the neuroplasticity of implicit motor sequence learning and the effect of acute exercise priming. Healthy adults (39.5% female) aged 22.55 ± 2.69 years were allocated to either a high-intensity exercise (HIIT) group (n = 16) or to a very low-intensity control group (n = 17). Following exercise, participants performed a serial reaction time task. MR spectroscopy estimates of sensorimotor gamma-aminobutyric acid (GABA) were acquired before and after exercise, and during task performance, and resting-state fMRI was acquired at the end of the protocol. We show that early stages of learning are linked to default mode network connectivity, while the overall degree of learning following sustained practice is associated with motor network connectivity. Sensorimotor GABA concentration was linked to the early stages of learning, and GABA concentration was modulated following HIIT, although the two were not related. Overall, via integration of multiple neuroimaging modalities we demonstrate that interactions between hippocampal and motor networks underlie *implicit* motor sequence learning.

**Key points summary:**

- Motor learning occurs across different temporal scales and can arise implicitly in the absence of conscious awareness.
- *Explicit* motor learning is linked to the brain’s primary inhibitory neurotransmitter, GABA, and interactions across motor and hippocampal networks.
- Whether these same neural mechanisms are implicated in *implicit* learning is unclear. Similarly, the capacity to influence learning via priming with cardiovascular exercise is yet to be clearly established.
- We show that early implicit learning is underpinned by default mode network connectivity and sensorimotor GABA concentration, while total learning following sustained practice is linked to motor network connectivity. We also found that HIIT exercise elevated sensorimotor GABA concentration, but not the magnitude of implicit learning.
- Overall, our results highlight shared involvement of default mode and motor networks in implicit motor sequence learning.

## Introduction

Motor skills are necessary to perform effectively and efficiently in our everyday lives. Acquiring a new skill (e.g., learning how to operate machinery or play a musical instrument) involves practicing a sequence of motor actions, with their execution becoming faster and more accurate with repetition and the passage of time. Research on the neural mechanisms supporting motor sequence acquisition has established a central role of the brain’s primary inhibitory neurotransmitter gamma-aminobutyric acid (GABA) (Coxon et al., 2014; Stagg et al., 2011a). Modulation of the GABA system is a prerequisite for the synaptic plasticity underlying learning and memory (i.e., long-term potentiation) (Bachtiar & Stagg, 2014), and is likely a factor contributing to inter-individual differences in learning capacity (Stagg et al., 2011a). GABA concentration within primary sensorimotor cortex (SM) is reduced as a motor sequence is acquired (Floyer-Lea et al., 2006; Kolasinski et al., 2019), and is associated with the degree of learning observed following practice (Kolasinski et al., 2019; Stagg et al., 2011a). SM GABA concentration is also linked to the strength of functional connectivity within the motor network (Stagg et al., 2014), indicating that interactions between the GABAergic system and task-relevant brain networks may underlie the learning of novel motor sequences.

Motor sequence learning occurs across multiple temporal scales, from acquisition during practice to offline consolidation over the minutes to hours and days that follow (Dayan & Cohen, 2011). The consolidation of motor skills over the minutes to hours following practice has been linked to modulation of GABA during acquisition (Stavrinos & Coxon, 2017), and to increased functional connectivity (Albert et al., 2009; King et al., 2022). Remarkably, motor consolidation might begin faster than previously thought, over the timescale of seconds, as evidenced by offline gains during short breaks between initial practice attempts (i.e., micro-consolidation) (Bönstrup et al., 2019, 2020). Evidence indicates that the micro-consolidation of an explicit motor sequence is mediated by activity across networks involving entorhinal and hippocampal regions (Buch et al., 2021; Jacobacci et al., 2020), and is also linked to beta oscillations across frontal-parietal regions (Bönstrup et al., 2019). Collectively, this evidence may reflect involvement of a distributed network involving frontal, parietal, and hippocampal regions including the default mode network, which has been proposed to mediate neural replay and consolidation of new memories (Kaefer et al., 2022).

Motor sequence learning can be explicit, investigated using tasks that directly cue the required motor sequence (as per aforementioned studies by Bönstrup et al., 2019, 2020; Jacobacci et al., 2020; Kolasinski et al., 2019), or implicit, where sequence knowledge is developed through repetition without explicit knowledge of the sequence (Robertson, Pascual-Leone, & Miall, 2004; Robertson, Pascual-Leone, & Press, 2004). Our recent work has established that offline micro-consolidation extends to implicit motor sequence learning (Brooks et al., 2024). However, the neural mechanisms linking micro-consolidation of implicit motor sequences to overall learning outcomes (following more sustained practice) are yet to be determined. Further, it remains unclear whether both motor and default mode networks are involved in the learning of *implicit* motor sequences. Addressing these gaps in knowledge will provide a more comprehensive understanding of how the brain encodes new motor skills across different temporal scales. Advancing knowledge in this regard will likely have implications for optimising human performance across a range of settings, such as in sporting contexts, and/or in the implementation of therapies aimed at rehabilitating motor skills following stroke and brain injury.

A considerable body of prior research has demonstrated that cardiovascular exercise primes neuroplasticity mechanisms implicated in learning and memory (Hendrikse et al., 2017). For example, regular exercise is associated with elevated markers of neuronal integrity within the hippocampus (Hendrikse et al., 2022), and a single bout of high-intensity interval exercise (HIIT) alters GABA concentration within non-exercised sensorimotor cortex (Coxon et al., 2018). Performing HIIT immediately prior to motor practice has been demonstrated to improve sequence consolidation (Roig et al., 2012; Stavrinos & Coxon, 2017; Taylor et al., 2024). However, these interactions may vary as a function of task demands and the time window under which they are assessed (Roig et al., 2013, 2016). The effect of HIIT exercise priming on neural mechanisms underpinning implicit motor sequence learning (i.e. modulation of GABA concentration and functional connectivity) remains unclear.

In summary, previous work has established a role of both hippocampal and motor systems in explicit motor sequence learning. The formation of novel motor skills is also linked to the activity of GABA, particularly within primary sensorimotor regions (Kolasinski et al., 2019; Stagg et al., 2011a). However, the effect of cardiovascular exercise on the neural mechanisms underlying sequence learning is yet to be clearly elucidated. Further, our understanding of how these neural mechanisms mediate different stages of *implicit* motor sequence learning (e.g., at the micro-scale vs following sustained practice) remains limited. Therefore, in this study we investigated the neuroplasticity of the initial learning of an implicit motor sequence, using MR spectroscopy and resting-state fMRI. High-intensity exercise was administered to prime task-relevant plasticity mechanisms in one group and compared to a very low-intensity control group. We expected that high-intensity exercise priming would modulate sensorimotor GABA concentration and facilitate motor sequence learning. We also hypothesised that early micro-offline consolidation of an implicit motor sequence would be associated with the coordinated activity of resting-state networks involving the hippocampus, i.e., functional connectivity of default mode network (as per Bönstrup et al., 2019; Jacobacci et al., 2020), but not domain-general networks (i.e., salience and visual networks). Lastly, we hypothesised that latter stages of motor skill learning (i.e., following more sustained practice, characterised by a reduced slope of the learning curve) would be associated with motor network connectivity and GABA concentration within sensorimotor cortex, and that this may be more pronounced in the HIIT group.

## Methods

### Ethical approval

This study was approved by the Monash University Human Research Ethics Committee (project # 27724) and all participants provided their written informed consent. The study conformed to the standards set by the Declaration of Helsinki, except for registration in a database. Participants were remunerated $30 for their participation.

### Participants

Thirty-eight healthy adults (39.5% female) aged 22.55 ± 2.69 years (mean ± SD; range 19 – 28) were recruited into the study. Participants reported no contraindications to magnetic resonance imaging or cardiovascular exercise, current prescription of psychoactive medication, nor history of psychological or neurological disorders. All participants were required to complete the long format of the International Physical Activity Questionnaire (IPAQ) prior to participation to assess self-reported physical activity. Handedness was determined by the Edinburgh Handedness Inventory. All participants were right-hand dominant (*M* = +90.42, *SD* = 23.72). Data from five participants were excluded due to developing explicit awareness of the repeating motor sequence during completion of the serial reaction time task (n = 4, see motor sequence learning for further detail), and an error in the MRI console software (n = 1). Thus, analyses were conducted with N = 33 (HIIT group, n = 16; control group, n = 17).

### Experimental design

A mixed experimental design was used to investigate interactions between exercise, implicit motor sequence learning, and MRI measures. Participants completed a 20-minute acute bout of either high-intensity interval cycling (HIIT group, n = 16) or a matched period of very low intensity cycling (Control group, n = 17). Following this, participants performed a serial reaction time task. MR spectroscopy estimates of GABA concentration were acquired at rest before and after exercise, and during performance of two runs of a serial reaction time motor task (Figure 1). Resting-state fMRI was also acquired at tbe end of the protocol to assess associations between network connectivity and learning. Self-report measures relating to exercise tolerance (Ekkekakis et al., 2005) and lifestyle factors that impact learning and memory (e.g., sleep quality) were also assessed (Buysse et al., 1989).

**Figure 1.**
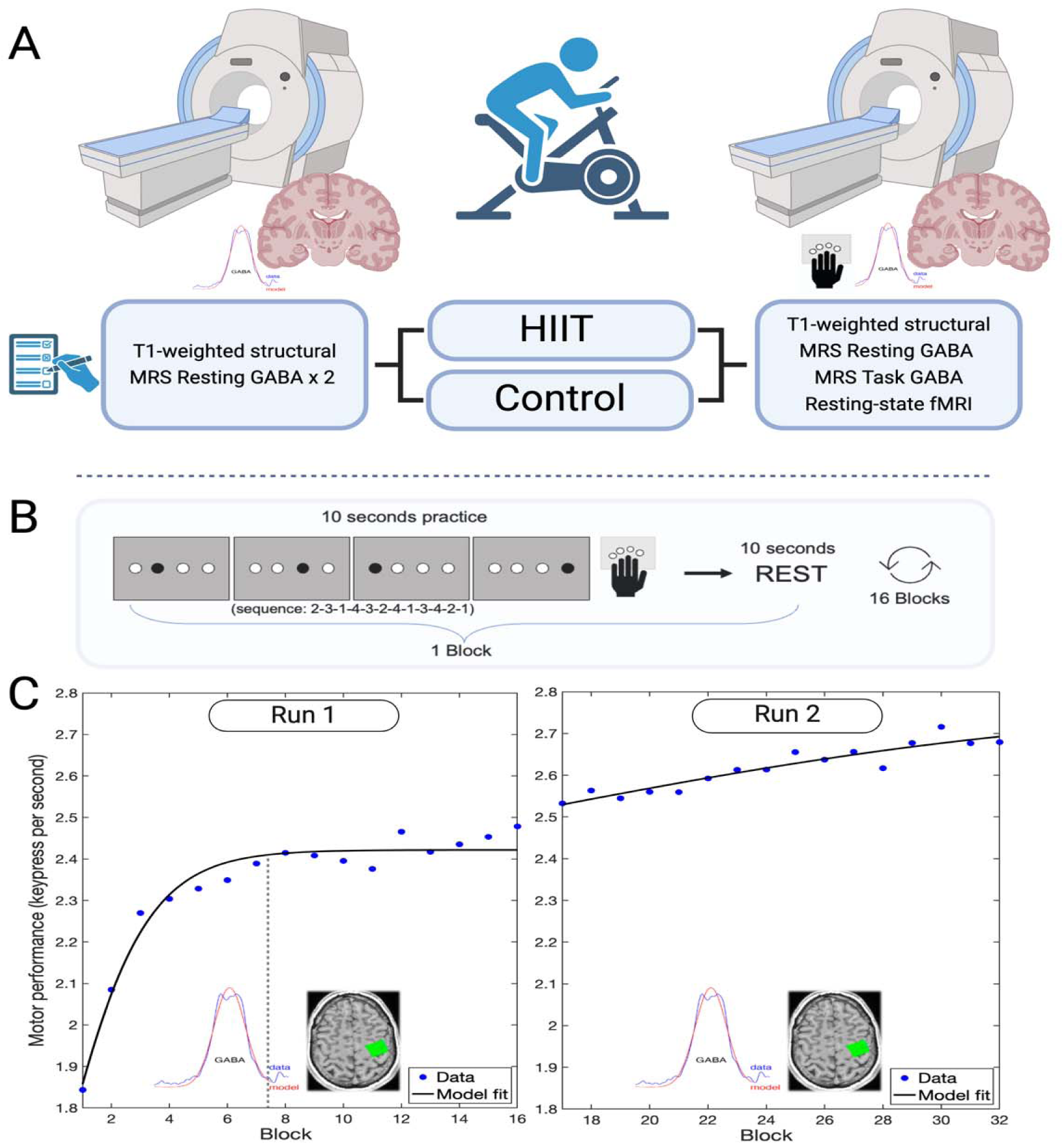
**(A)** Experimental overview. A mixed experimental design was used to investigate associations between exercise, motor learning, and MRI measures. Participants completed a 20-minute acute bout of either high-intensity interval cycling (HIIT group, n = 16) or a matched period of very low intensity cycling (control group, n = 17). **(B)** Following this, participants performed two runs of a serial reaction time task which included an implicit 12-item repeating sequence. Each run comprised 16 blocks of 10 second periods of practice and rest. MR spectroscopy estimates of GABA+ concentration were acquired before and after exercise, and during performance of the motor task. Resting-state fMRI was also acquired to assess associations between network connectivity and learning. **(C)** Motor task performance is represented as the mean increase in correct keypresses per second across blocks for all participants (blue circles), modelled with an exponential function, B(t) ∼ L(t) = k1 + (k2 / 1 + e^−k3t^), where k1 and k2 determine learning plateau, k3 the steepness of the learning function, and t ∈ [1, +∞) represents trial (t) starting from 1 and extending to infinity (black line, Bönstrup et al., 2019). The model was fit separately to each block to delineate the ‘early learning’ phase cut-off, by which point 95% of early learning had occurred (Bönstrup et al., 2019). This is represented by the vertical dotted line. GABA+ concentration was measured across both runs of the motor task.

### Exercise

Participants were pseudo-randomised into HIIT or control groups using an algorithm that minimised variance between groups with regard to biological sex and age. Exercise intensity was tailored to each individual based on their heart rate reserve (HRR), where: HRR = Age predicted maximum heart rate – resting heart rate (Tanaka et al., 2001). Heart rate was continuously monitored throughout the exercise session using a Polar H7 monitor (Polar Electro, Kempele, Finland). HIIT exercise involved cycling on a stationary cycle ergometer (Wattbike, Geelong, Australia) at alternating epochs of moderate intensity (3-minutes, target heart rate of 50-60% HRR) and high intensity (2-minutes, target heart rate of 90% HRR for the final epoch) for a total duration of 20-minutes, as per our previous work (Cadwallader et al., 2024; Coxon et al., 2018; Curtin et al., 2023; Stavrinos & Coxon, 2017; Taylor et al., 2024). The control group were required to turn the pedals at a very low cadence (target heart rate was to remain below 20% HRR) requiring minimal exertion. Here, we were interested in assessing the systemic effects of HIIT exercise on motor plasticity. As such, in line with previous studies by Coxon et al., 2018, we utilised lower limb-based stationary cycling and recorded GABA concentration from the non-exercised hand representation in primary motor cortex.

### Motor sequence learning – serial reaction time task

To assess motor sequence learning, we utilised a customised version of the serial reaction time task (SRTT). The SRTT was commenced approximately 20-25 minutes following exercise (Figure 1). Participants lay supine in the scanner and viewed four white circles arranged horizontally on a monitor screen, with their non-dominant (left) hand positioned over a four-button box. Their little finger aligned to the left most button (and left most circle #1) and index finger to the right most button (and right most circle #4). Participants were instructed to respond to the circle that turned black by pressing the corresponding button as quickly and accurately as possible. Undisclosed to participants, the visual cues followed a 12-item repeating sequence (2-3-1-4-3-2-4-1-3-4-2-1) (Figure 1B). A correct response was required to progress to the next cue. The task alternated every 10-seconds between task practice and rest for a total of 16 practice blocks, 320-seconds in total. Participants completed two separate runs of the task (i.e. a total of 32 practice blocks, 640-seconds) to assess the initial learning of an implicit motor sequence, and associated changes in GABA concentration.

Following completion of the session, participants were questioned about their level of explicit sequence awareness. Participants who were able to accurately replicate the first five or more sequence items were determined to have gained some degree of explicit sequence awareness (HIIT, n = 2, Control, n = 2), and were removed from subsequent analyses (Brooks et al., 2024).

### Magnetic resonance imaging (MRI)

MRI data were acquired with a Siemens 3T Skyra scanner and 32-channel head coil. T1-weighted structural images (Magnetization Prepared Rapid Gradient Echo, TR = 2.3 s, TE = 2.07 ms, voxel size 1mm^3^, flip angle 9°, 192 slices) and single voxel GABA-edited Mescher-Garwood Point Resolved (MEGA-PRESS) spectra were acquired at baseline prior to exercise. T1-weighted structural data was inspected to localise each participant’s hand knob region of the central sulcus. MEGA-PRESS data were acquired from a 3 x 3 x 3 cm voxel positioned in sensorimotor cortex (SM), centred over the hand knob in the right hemisphere (i.e., contralateral hemisphere to left hand used to complete the motor task). Two assessments of resting GABA concentration were acquired prior to exercise (TE = 68 ms, TR = 1.5 s, edit pulse bandwidth = 45 Hz, editing pulses at 1.9 ppm and 7.5 ppm, 105 ON-OFF averages at each acquisition). Repeat assessments of resting-state GABA were conducted to establish the stability of MRS measures prior to exercise, as per our previous work (Coxon et al., 2018). A reference spectrum, comprising 16 ON-OFF averages with unsuppressed water was also obtained using the same parameters.

Following exercise, a second T1-weighted image was acquired to inform accurate voxel placement relative to the baseline assessment (see Figure 2, voxel overlap). SM GABA concentration was assessed again at rest (105 averages), and across both runs of the SRTT (105 averages each run, 210 averages total). Resting-state Echo Planar Images (TR = 2.46 s, TE = 30 ms, voxel size 3mm^3^, flip angle 90°, 44 slices, 7.34 mins duration) with whole brain coverage were acquired at the end of the protocol. This was acquired at the end of the protocol to avoid any impact of gradient coil heating from the echo planar imaging sequence on magnetic field homogeneity and the stability of MR spectroscopy estimates (Hui et al., 2021). During the resting-state scan, participants were instructed to keep their eyes open and focus on a black fixation cross presented on a white background, whilst trying not to think of anything in particular.

**Figure 2.**
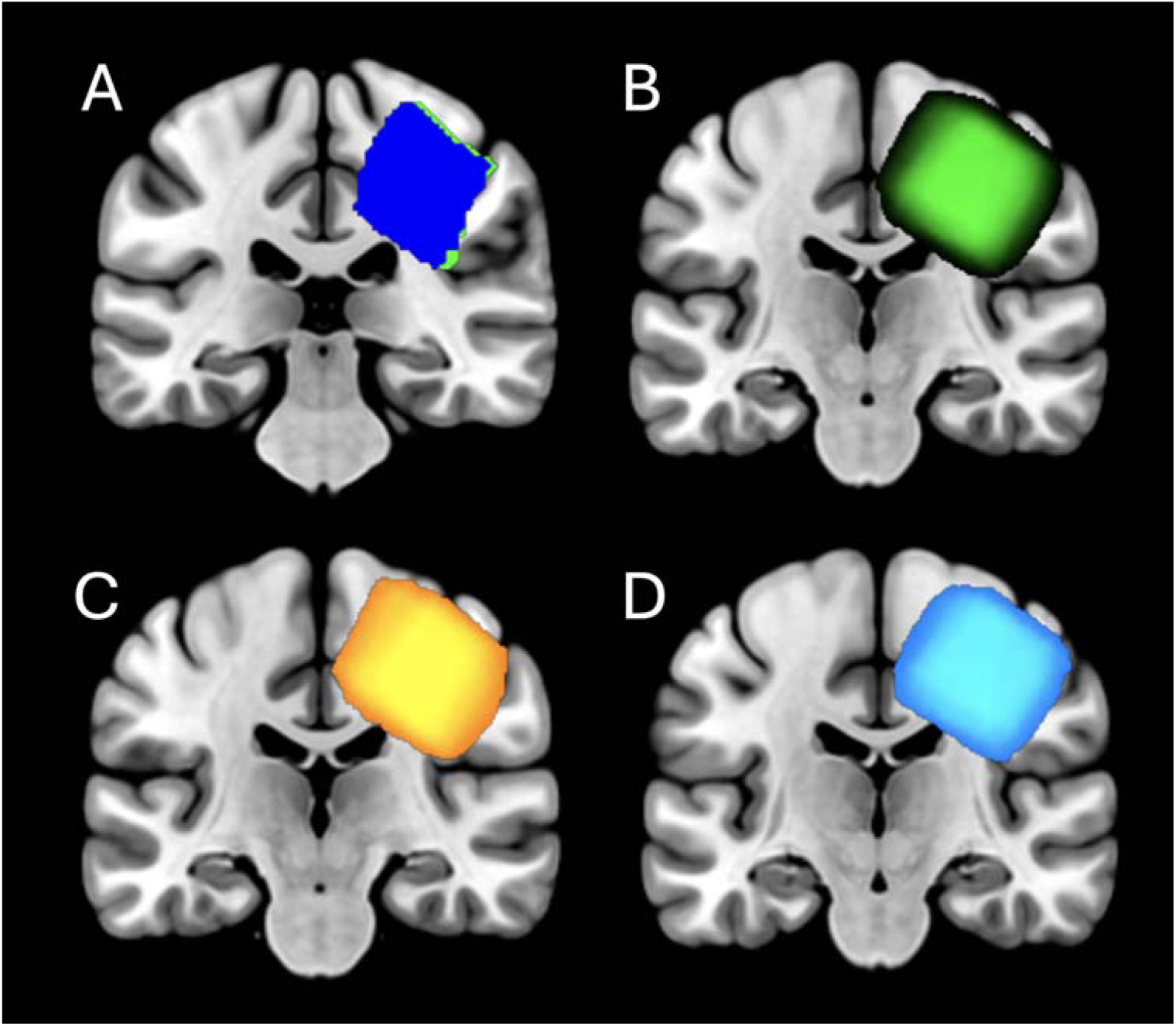
Positioning of magnetic resonance spectroscopy voxel centred over hand region of sensorimotor cortex. MEGA-PRESS data were acquired from a 3 x 3 x 3 cm voxel before and after exercise, and during performance of a serial reaction time task. The overlap in voxel position pre-post exercise is shown for a representative participant **(A)**, all participants (N = 33) **(B)**, and separately for HIIT (n = 16) **(C)** and control (n = 17) **(D)** groups, where greater spatial overlap is represented by lighter coloration (B-D).

### Data analysis

#### Motor sequence learning

As per previous work (Bönstrup et al., 2019; Brooks et al., 2024), motor skill performance was conceptualised as number of correct responses per second. The learning trajectory was modelled at the group-level (N = 33) using an exponential function (Bönstrup et al., 2019; Brooks et al., 2024, Figure 1C).

Motor learning was assessed at the micro-scale (i.e., micro-consolidation) according to our previous work (Brooks et al., 2024). In brief, we investigated rapid improvements in performance during both active practice (i.e., micro-online gains) and the brief 10 second periods of rest (i.e., micro-offline gains). To assess micro-online and -offline changes, analysis was restricted to an ‘early learning’ phase, where 95% of learning occurred (i.e., blocks 1-7 on the first run of the task, Figure 1C) (Brooks et al., 2024). A sliding average window of four correct responses was used, and the mean tapping speed (quantified as keypresses/s) across the first and last four correct key presses in each block represented performance at the start and end of each block, respectively. Micro-offline changes were quantified as the summed change between blocks, i.e., the summed change in tapping speed between the start of block n+1 and the end of block n. Micro-online changes were the summed change within blocks, i.e., the summed change in tapping speed between start and end of block n. Early learning was calculated by summing micro-online and micro-offline values. We also assessed the total degree of learning across performance of both task runs. Total learning was defined as the change in tapping speed from the first four correct keypresses of block 1 to the last four correct keypresses of block 32 (i.e., end of second run).

#### Quantification of GABA

Gannet (version 3.2.0) (Edden et al., 2014) was used to analyse MEGA-PRESS spectra. Estimates of GABA concentration (3.0 ppm) were calculated relative to the unsuppressed water signal from the voxel centred over right SM. Pre-processing of MRS data (GannetLoad module), included steps for phased-array channel combination of raw Siemens ‘twix’ files, exponential line broadening (3 Hz), Fourier transformation to yield time-resolved frequency-domain spectra, frequency and phase correction, outlier rejection, and time averaging of the edited difference spectrum. The area of the edited GABA peak at 3 ppm was quantified using a single Gaussian peak model, and a nonlinear least-squares fitting approach (GannetFit module). To acknowledge a possible macromolecule contribution to the edited spectra, GABA+ is referred to in place of GABA for all relevant analyses (Edden et al., 2014). As an internal reference, GABA+ concentrations are expressed in institutional units, relative to the unsuppressed water signal estimated with a Gaussian-Lorentzian model.

Resting GABA was quantified at baseline prior to exercise, and again following exercise (HIIT or Control condition). GABA was also quantified during different phases of motor skill learning. To yield an estimate of GABA during the ‘early learning’ phase, the mean of the spectra acquired during the first 52 MEGA-PRESS on-off acquisitions was calculated (i.e., the midpoint of the MRS sequence encompassing blocks 1-8 of the SRTT). To examine change in GABA concentration across the SRTT task, early learning GABA was subtracted from late learning GABA (the last 52 MEGA-PRESS on-off acquisitions, corresponding to blocks 24-32 of the SRTT).

Partial volume correction was also performed. The MRS voxel was co-registered to each subject’s anatomical T1-weighted image using the GannetCoRegister module, and partial volume segmentation of the T1-weighted anatomical image within the SM voxel was then conducted using GannetSegment. Partial volume correction of GABA concentration was then performed by removing cerebrospinal fluid fractions (GannetQuantify) to yield corrected GABA+/H_2_O ratios. The full width at half maximum of GABA signals (FWHM) and GABA signal fit error (relative to the unsuppressed water peak, EGABA) were used to assess the quality of GABA+ model fit. Spectra with an EGABA value of *>*20% or exceeding a z-score of 3.29 were labelled as outliers and removed from the analysis. FSLmaths was used to calculate the percentage of voxel overlap between baseline and post-exercise assessments (as depicted in Figure 2).

#### Resting-state fMRI analysis

Data were pre-processed using fMRIprep (version 1.1.1), with default parameters including ICA-AROMA denoising and susceptibility-derived distortion estimation (AFNI 3dqwarp). This resulted in spatially realigned Echo Planar Images in normalised space (normalised to the MNI152NLin6Asym template). To examine associations between functional connectivity and motor performance, we then conducted a dual-regression analysis in FSL (Jenkinson et al., 2012), according to the approach outlined in Stagg et al., (2014). Briefly, each pre-processed resting-state fMRI scan was temporally concatenated across participants to yield a single 4D dataset. Data were then split into 25 components using independent component analysis. Resting-state networks of interest were identified via spatial comparison with previous defined networks (Beckmann et al., 2005). The motor and default mode network were the primary networks of interest given previous evidence implicating hippocampal and motor regions in motor sequence learning at both micro-(Buch et al., 2021; Jacobacci et al., 2020) and macro-scale (Dupont-Hadwen et al., 2019; Kolasinski et al., 2019). The salience network and visual network were also identified as control networks to assess anatomical specificity (see figure 3). The dual-regression analysis was then conducted to establish subject-specific functional networks. This analysis involves a two-stage regression sequence, in which a spatial regression was first performed against single-subject data using all 25 components, followed by a temporal regression of the resulting normalised time-course to generate subject-specific maps. This map was taken as a subject-specific estimate of functional connectivity. Binarised masks were then generated for each resting-state network of interest, derived from the group average of pre-processed 4D images (multiple comparison corrected, p < .05). The mean value was calculated for each masked network for each subject to yield estimates of average network connectivity.

**Figure 3.**
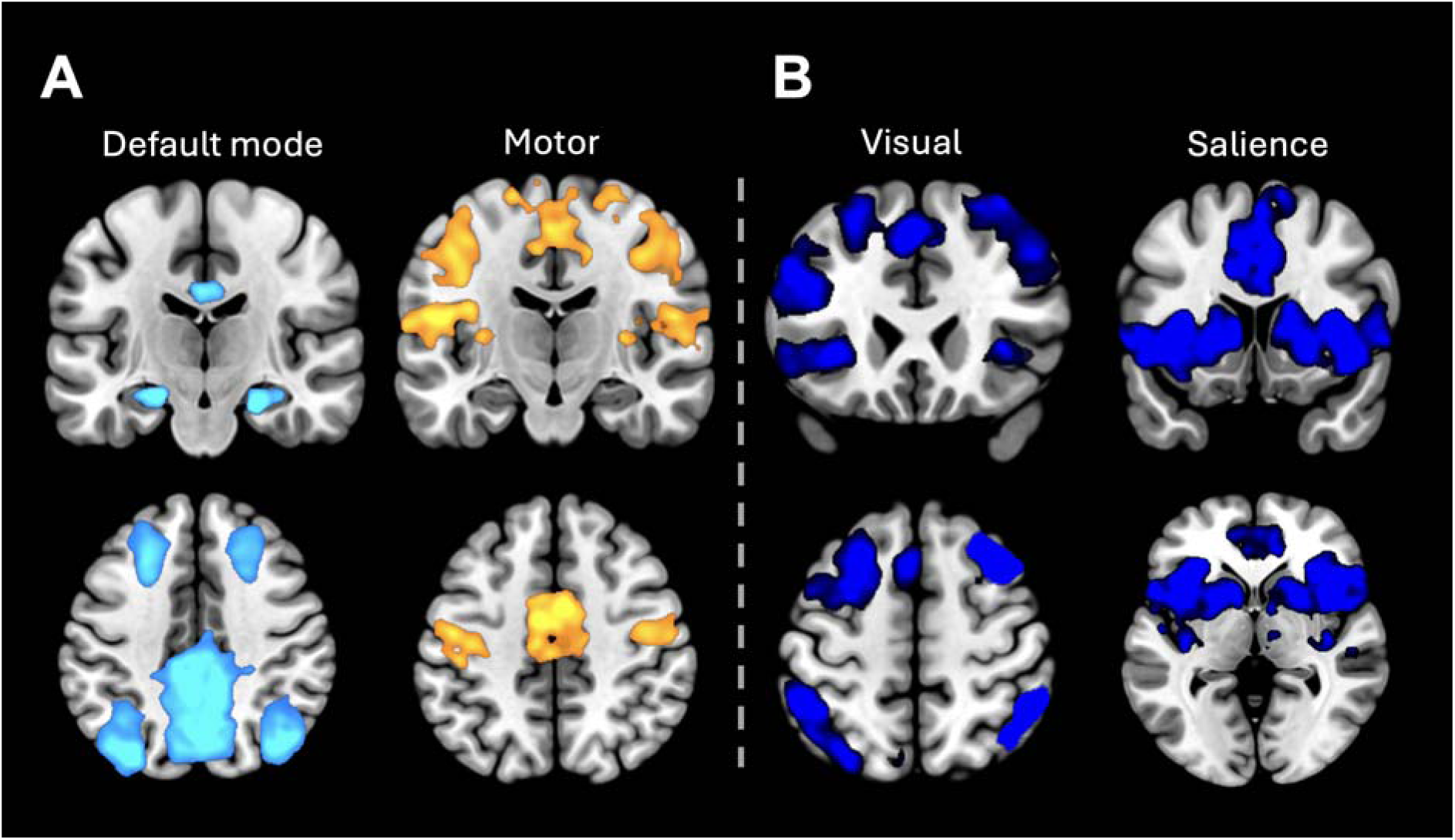
Visualisation of resting-state networks. Functional maps derived from all participants (N = 33) are shown for primary networks of interest **(A)** including default mode (light blue colouration) and motor networks (yellow-orange colouration), and control networks **(B)** including visual and salience networks (dark blue colouration).

#### Statistical approach

Statistical analysis was conducted using JASP (v 0.10.2). An alpha level of .05 was adopted for all inferential statistics conducted under the null hypothesis significance testing framework. To quantify the relative evidence for the alternative vs. the null hypothesis, equivalent Bayesian statistics were also determined (the prior was set in support of the alternative over the null hypothesis, i.e. BF_10_). For Bayesian tests, BF_10_ > 3 was considered strong evidence against the null, and BF_10_ < 0.33 as strong evidence for the null. For Bayesian t-tests, the Cauchy parameter was set to a conservative default value of 0.707 (Ly et al., 2016; Rouder et al., 2009).

### Comparisons between HIIT and Control conditions

To assess the effect of HIIT on measures of sequence learning and MRI measures (i.e., GABA, functional connectivity), independent-samples t-tests were conducted between HIIT and control groups. In the case of violations of normality, the nonparametric equivalent (i.e. Mann-Whitney U test) was used. In our previous work, we showed that a single dose of HIIT did not significantly impact consolidation at the micro-scale (Brooks et al., 2024). Therefore, here, we restrict our group comparisons to imaging measures and total learning outcomes. To account for differences in baseline GABA+ concentration, difference scores were calculated by subtracting baseline GABA+ estimates from post timepoint (i.e., post exercise – baseline).

### Relationship between motor plasticity and learning

Based on past evidence linking hippocampal dynamics to micro-offline gains in explicit learning (Buch et al., 2021; Jacobacci et al., 2020), bivariate correlations were conducted to investigate associations between resting-state functional connectivity and micro-offline consolidation of an implicit motor sequence (data pooled across exercise groups, bonferroni-adjusted for the number of included networks, α-level = .013). Multiple linear regression models were conducted to examine how micro-offline consolidation and total learning related to functional connectivity and change in GABA concentration measured across motor practice. Separate models were conducted on learning measures at the micro and overall (macro) scale.

## Results

Exercise was well tolerated by participants and no adverse events were reported following HIIT. There were no significant differences between HIIT and Control groups in terms of age (p = .68), biological sex (p = .75), pre-existing physical activity (p = .30), or sleep quality (p = .21) (see Table 1).

**Table 1.** Comparison of participant characteristics between HIIT and Control groups (Mean ± Standard Deviation).

|  | HIIT | Control | *p* value |
| --- | --- | --- | --- |
| N (% Female) | 19 (36.84%) | 19 (42.11%) | .75 |
| Age (years) | 22.37 ± 2.65 | 22.74 ± 2.79 | .68 |
| BMI (kg/m<sup>2</sup>) | 22.06 ± 2.21 | 23.3 ± 4.14 | .26 |
| IPAQ (MET) | 4800 ± 3220 | 3783 ± 2739 | .30 |
| PRETIE-Q T | 26 ± 5.08 | 23.95 ± 4.92 | .21 |
| PRETIE-Q P | 26.32 ± 5.09 | 23.37 ± 4.22 | .06 |
| PSQI | 5.05 ± 2.53 | 4.16 ± 1.77 | .21 |
| RHR (bpm) | 64.26 ± 9.75 | 65.37 ± 10.07 | .92 |
| Exercise HR (bpm) | 180.68 ± 7.10 | 89.95 ± 10.96 | **<.001** |
| %HRR | 93.9 ± 3.3 | 49.6 ± 13.3 | **<.001** |

*Note.* BMI: Body Mass Index. IPAQ: International Physical Activity Questionnaire (METS, metabolic equivalent minutes per week), higher values indicate greater physical activity levels. PSQI: Pittsburgh Sleep Quality Index global score, with higher values indicating poorer sleep quality. PRETIE-Q: Preference for and Tolerance of the Intensity of Exercise Questionnaire, higher exercise intensity tolerance (PRETIE-Q T) and preference (PRETIE-Q P) are indicated by higher values. RHR: Resting Heart Rate (seated), bpm = beats per minute. For Exercise Heart Rate the peak of the last minute of exercise is reported for HIIT, and the 20-minute average is reported for the low intensity Control group. %HRR: percentage of heart rate reserve achieved during exercise, where HRR = age-predicted heart rate max – resting heart rate).

### Effect of HIIT on GABA and learning

We first examined the effects of HIIT on resting SM GABA+ concentration. Two assessments of GABA+ were acquired prior to exercise to verify stability and reliability of the measure. No significant differences were observed between GABA+ baseline assessments when compared across the entire sample [t(1,32) = 0.66, p = .52, Cohen’s d = 0.11, BF_10_ = 0.23], nor within the HIIT [t(1,15) = 1.50, p = .15, Cohen’s d = 0.38, BF_10_ = 0.65] or active control [t(1,16) = −0.53, p = .61, Cohen’s d = 0.13, BF_10_ = 0.28] groups separately. Therefore, the average of the two baseline GABA+ assessments was calculated and used for subsequent analysis. No significant differences in baseline SM GABA+ concentration were observed between HIIT and control groups [t(1,31) =-0.96, p = .35, Cohen’s d = 0.33, BF_10_ = 0.47], indicating that groups were appropriately matched. There were also no differences in the quality of the GABA+ model fit (EGABA) or the unsuppressed water signal (used as a reference for GABA+ quantification) between groups at baseline [water: t(1,31) =-0.47, p = .65, Cohen’s d = 0.16, BF_10_ = 0.36; EGABA: t(1,31) =0.33, p = .74, Cohen’s d = 0.12, BF_10_ = 0.35], nor following exercise [water: t(1,31) =-0.78, p = .44, Cohen’s d = 0.27, BF_10_ = 0.42; EGABA: t(1,31) =-1.05, p = .30, Cohen’s d = 0.37, BF_10_ = 0.51]. To examine changes in GABA following exercise, baseline concentration was subtracted from post assessment (Δ GABA = GABA post – GABA pre). In line with our previous findings (Coxon et al., 2018), we observed modulation of resting SM GABA+ concentration following a bout of HIIT. A significantly greater increase in GABA concentration was observed following HIIT exercise, relative to the control group [t(1,31) = 2.23, p = .033, Cohen’s d = 0.78, BF_10_ = 2.09].

Despite the exercise-related change in GABA+, the groups did not differ behaviourally in terms of their learning on the SRTT. Specifically, there were no differences between HIIT and control groups for early learning [Task Run 1: t(1,31) = 0.18, p = .86, Cohen’s d = 0.06, BF_10_ = 0.34] or total learning [Across runs: t(1,31) = 0.29, p = .77, Cohen’s d = 0.10, BF_10_ = 0.34] (Figure 4). Similarly, there were no differences observed in GABA+ concentration between groups during motor task performance [all timepoints p > .05]. Overall, we provide evidence that HIIT modulates resting SM GABA+ but not implicit motor sequence learning at the microscale (Brooks et al., 2024), or when total learning across this task is considered.

**Figure 4.**
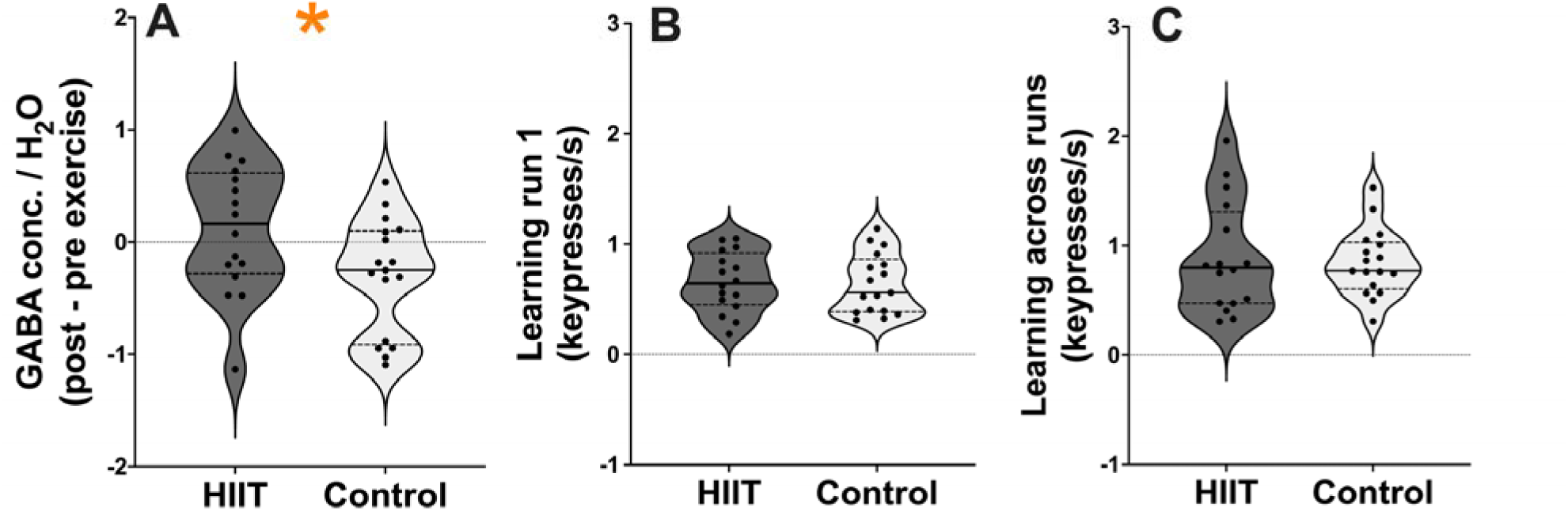
Comparison of SM GABA+ concentration and motor learning between HIIT and Control groups. **(A)** Comparison of GABA concentration between groups. **(B)** Comparison of learning across the first practice run of the SRTT between groups. **(C)** Comparison of learning across both runs of the SRTT between groups. For all plots, filled circles indicate individual participants. For each exercise group, median values are indicated by the horizontal bolded line, and higher and lower quartiles are indicated by dashed lines. * p < .05.

### Micro-offline consolidation and default mode network connectivity

Previous evidence has linked the micro-consolidation of an explicit motor sequence to the activity of networks involving the hippocampus (Buch et al., 2021; Jacobacci et al., 2020). In our recent work, we demonstrate that micro-consolidation generalises to implicit motor sequence learning (Brooks et al., 2024). Here, we were interested in whether micro-consolidation of an implicit motor sequence was also associated with default mode network connectivity, in which the hippocampus is a primary node. We observed a significant positive association between micro-offline gains and resting-state connectivity in the default mode network (r = .44, p_Bonf_ = .044, BF_10_ = 4.73). This relationship was specific to the default mode network; no other resting-state networks showed significant associations with micro-offline gains (Motor: r = .35, p_Bonf_ = .24, BF_10_ = 1.29, Salience: r = .24, p_Bonf_ = .68, BF_10_ = 0.53; Visual: r = .22, p_Bonf_ = .88, BF_10_ = 0.45) (Figure 5). In summary, default mode network activity is associated with the rapid micro-consolidation of implicit motor sequences occurring during brief periods of rest.

**Figure 5.**
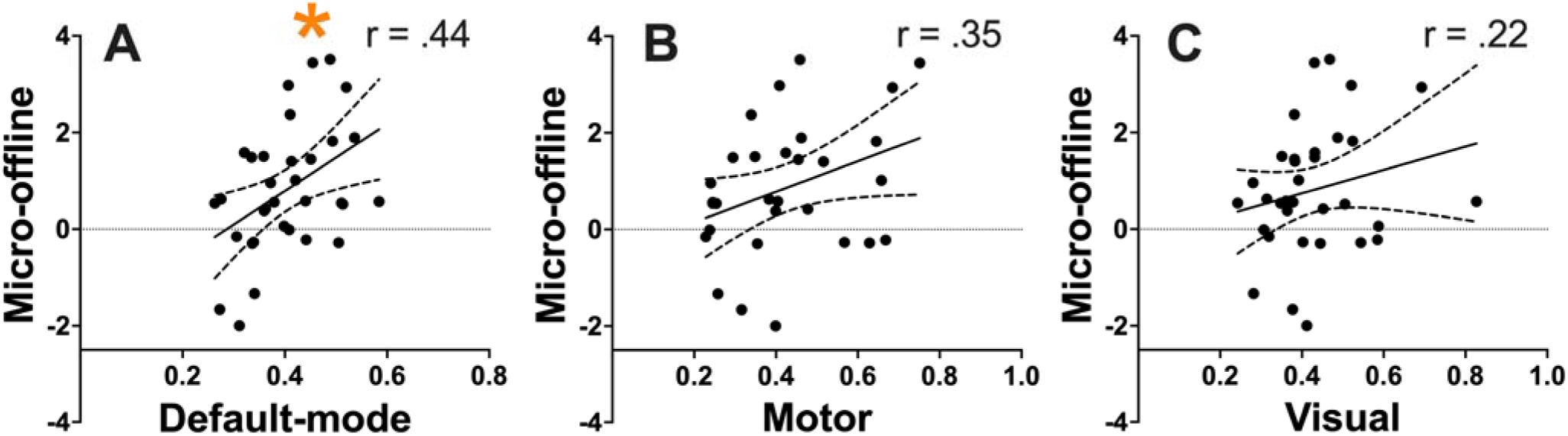
Associations between micro-offline consolidation and resting-state functional connectivity. **(A)** Significant positive correlation between micro-offline gains and default mode network connectivity. No other significant correlations were observed between micro-offline consolidation and other resting-state networks **(B)** Motor network, and **(C)** Visual network. Circles represent individual values, solid line shows regression line of best fit, dashed line depicts 95% confidence interval. Micro-offline gains in implicit sequence learning are represented on the y-axis and averaged functional network connectivity is represented on the x-axis. * p < .05.

### Motor sequence learning, GABA concentration, and functional connectivity

GABA regulates the plasticity underlying motor learning (Stagg et al., 2011a), with change in SM GABA concentration associated with motor sequence learning magnitude (Kolasinski et al., 2019). Here, we extend on these findings, and examine how resting-state network connectivity, GABA are involved across early and later stages of implicit sequence learning, and whether a single dose of HIIT exercise influences these relationships.

We first assessed the relationship between early learning, GABA+, and hippocampal network connectivity. GABA+ measured during early learning and resting-state default mode connectivity explained a significant proportion of the variance in early sequence learning [Model summary: R^2^ = .27, Adjusted R^2^ = .19, F(3,32) = 3.51, p = .028]. We observed a negative correlation between early learning and GABA+ [B = −0.14 (−0.26, −0.03 95% CI), p = .015, 13% unique variance explained], and a positive correlation between early learning and default mode connectivity [B = 0.709 (0.12, 1.51 95% CI), p = .021, 7% unique variance explained]. Notably, these associations were specific to GABA+ measured during early learning and default mode connectivity; no other resting-state connectivity measure nor estimates of GABA+ across other timepoints explained a significant proportion of variance in early motor sequence learning [all p > .05]. Exercise condition (HIIT vs Control) was not a significant predictor in the model [B = −0.09 (−0.21, 0.06 95% CI), p = .248, 3% unique variance explained]. Overall, we provide evidence that default mode network connectivity and GABA+ measured during learning support early motor skill consolidation.

We next investigated the neurophysiological correlates of sequence learning across the entire practice period, i.e. total learning across task blocks 1-32. Two motor connectivity estimates were identified as outliers (z-score > 2.7) and removed from the analysis. Previous work has linked sequence learning across similar timespans to that seen here with motor network plasticity (Pascual-Leone et al., 1994; Tzvi et al., 2014). Indeed, a regression model with motor network connectivity and Δ GABA+ explained a significant proportion of the variance in total sequence learning [Model summary: R^2^ = .27, Adjusted R^2^ = .19, F(3,28) = 3.13, p = .044]. We observed a positive correlation between total learning and motor network connectivity [B = 1.59 (0.63, 3.07 95% CI), p = .002, 18% unique variance explained]. This relationship was specific to motor connectivity; no other resting-state network explained a significant proportion of variance in total learning when substituted or added as an additional predictor in the model [all p > .05]. Non-significant associations were observed between total learning and Δ GABA+ [B = 0.11 (−0.10, 0.37 95% CI), p = .29, 3% unique variance explained], and total learning and exercise condition [B = 0.02 (−0.43, 0.28 95% CI), p = .92, 0% unique variance explained]. Overall, we provide evidence implicating GABA and both motor and default mode network connectivity in the consolidation of an implicit motor sequence.

## Discussion

In this study, we investigated the neural mechanisms supporting implicit motor sequence learning, and the influence of high-intensity cardiorespiratory exercise (HIIT) priming. Previous evidence has established a central role of sensorimotor GABA in motor learning (Kolasinski et al., 2019; Stagg et al., 2011a), and involvement of hippocampal and motor networks in *explicit* motor sequence learning at both the micro-scale (Buch et al., 2021; Jacobacci et al., 2020) and following more sustained practice (Stagg et al., 2011a). Here, we report evidence that these systems support the learning of an *implicit* motor sequence. We show that micro-offline gains in implicit sequence learning are linked to default mode connectivity. Further, we demonstrate that the early stages of implicit sequence learning are linked to default mode connectivity and sensorimotor GABA concentration, and that total learning following more sustained practice is associated with the functional connectivity of the motor network. Our results indicate that interactions between default mode and motor networks underlie initial *implicit* motor learning. Investigation of the priming effects of HIIT exercise replicated our previous study (Coxon et al., 2018) showing sensorimotor GABA is modulated by HIIT. However, in contrast to our hypothesis, the priming effect of HIIT on sensorimotor GABA did not influence the magnitude of sequence learning.

### The effect of high-intensity interval training (HIIT) on MRI measures of motor plasticity

A growing body of evidence indicates that cardiorespiratory exercise impacts GABAergic neurotransmission (Hendrikse et al., 2017). For instance, increased GABA concentration has been reported in visual cortex following 8-17 minutes of moderate/high-intensity exercise (Maddock et al., 2016). Further, our previous work established that there is a consistent elevation in sensorimotor GABA concentration following a single session of HIIT (Coxon et al., 2018). Several studies have demonstrated modulation of GABA-mediated synaptic inhibition in motor cortex with transcranial magnetic stimulation measures following both moderate-(Singh et al., 2014) and high-intensity exercise (Cadwallader et al., 2024; Curtin et al., 2023). In the present study, we replicate our previous spectroscopy study (Coxon et al., 2018), demonstrating increased resting sensorimotor GABA concentration following a 20-minute dose of HIIT relative to a control condition. We show this in a larger sample than previously reported, establishing the effect is robust.

Modulation of GABA has been associated with the HIIT exercise-related increases in blood lactate concentration (Coxon et al., 2018). Systemic concentrations of lactate increase following high-intensity exercise (Brooks et al., 2005), and lactate is implicated in signalling pathways regulating the de novo synthesis of GABA in the brain (Maddock et al., 2016). The increase in GABA we observed may also relate to circulation of brain-derived neurotrophic factor, which is increased in response to high-intensity exercise (Saucedo Marquez et al., 2015), and involved in regulation of GABAergic neurotransmission (Cash et al., 2017). MRS provides an estimate of the overall tissue concentration of GABA, which on a functional level likely reflects extrasynaptic GABAergic tone (Stagg et al., 2011b). Hence, collectively these findings highlight the effect of high-intensity cardiovascular exercise on tonic GABA activity within sensorimotor cortex. Future work is required to delineate the specific physiological mechanisms mediating these effects.

In contrast to our hypotheses, modulation of sensorimotor GABA by HIIT-exercise priming did not influence implicit sequence learning. We observed no differences between HIIT and control conditions for the earliest micro-phase of learning (‘fast’ learning), nor when comparing the total magnitude of learning across more extended practice (approximately 11 minutes, total task duration). Similarly, no associations were observed between learning and exercise in the regression analyses. This result extends upon our recent work (Brooks et al., 2024), but conflicts with some previous work reporting facilitation of implicit sequence learning following exercise (Roig et al., 2013). For example, improved encoding (Mang et al., 2014) and retention (Mang et al., 2016) of implicit motor sequences have been demonstrated following a single dose of HIIT exercise. The reason for these inconsistent outcomes is unclear, though this may relate to differences in task demands across studies.

In the current study, we utilised a serial reaction time task as a measure of implicit sequence learning. We observed a rapid improvement in skill, with 95% of learning occurring by block 7 at the group level (i.e., following 140 seconds of practice). In comparison, studies reporting an effect of HIIT-priming on implicit learning have utilised more complex tasks with comparatively protracted learning trajectories (Mang et al., 2016). Plausibly, the additional benefit of HIIT priming on sequence learning may be more likely to manifest in the context of more challenging sequence learning paradigms, though future work is required to assess the validity of this interpretation.

An alternative explanation for these inconsistent outcomes across studies relates to the temporal coupling between exercise and learning. Previous studies reporting exercise-related facilitation of implicit learning have commenced motor skill practice immediately following exercise (Mang et al., 2014, 2016). In the current study, owing to the timing of our MRI protocol in which a T1-weighted structural image and resting estimates of GABA concentration were acquired prior to the task, motor practice was commenced ∼20-25 minutes after exercise cessation. Indeed, the temporal proximity of exercise to learning has been cited as a factor that may influence subsequent outcomes (Roig et al., 2016). In this sense, a closer temporal coupling of HIIT to motor learning may have maximised exercise-related benefits. However, on the other hand, both fatigue and high levels of physiological arousal (e.g., as commonly experienced immediately following HIIT) are also known to impact memory performance (Park, 2005), and may otherwise mask the cognitive benefits of exercise. For example, a single bout of moderate-high intensity exercise performed four hours after encoding improved retention on declarative memory tasks, whereas exercise immediately after encoding did not (van Dongen et al., 2016). This result indicates that exercise-related effects on learning may vary across time depending on physiological states, phase of memory formation, and the brain regions involved. Collectively, these diverging outcomes suggest a complex relationship between exercise and memory formation, likely influenced by task demands and the time window under which they are assessed.

### Default mode and motor systems support implicit sequence learning

In line with our expectations, we provide the first evidence that micro-offline consolidation of an implicit motor skill is linked to the activity of brain networks involving the hippocampus. We observed a significant positive correlation between default mode functional connectivity strength and individual micro-offline skill gain. This relationship was specific to the default mode network; no other relationships between offline skill gain and functional connectivity were observed across motor, salience, or visual networks. This finding extends on previous work which has linked micro-offline consolidation of explicit motor sequence learning to hippocampal dynamics, including neural replay bursts across a cortical-hippocampal network (Buch et al., 2021), and changes in both blood-oxygen level dependent activity and microstructure within hippocampus and precuneus (Jacobacci et al., 2020). The default mode network acts as a primary hub for neural replay (Higgins et al., 2021), and may mediate the encoding of novel experience within the hippocampus to longer-term consolidation across a distributed cortical representation (Kaefer et al., 2022). Our finding adds to a growing body of literature linking short-term consolidation of procedural motor memories during wakefulness to the coordinated activity of brain networks involving the hippocampus – a region classically associated with declarative learning taxonomies (Henke, 2010). Our finding is correlational, and further work is required to establish causal links between default mode network dynamics and the micro-consolidation phenomenon.

Converging lines of evidence indicate that explicit motor learning is underscored by interactions across both sensorimotor and hippocampal systems (Deleglise et al., 2023; Della-Maggiore, 2024; Döhring et al., 2017; Jacobacci et al., 2020). Explicit sequence learning has also been linked to dynamic changes between networks, with a decrease in sensorimotor GABA concentration during learning (Floyer-Lea et al., 2006; Kolasinski et al., 2019), and a shift towards decreased hippocampal connectivity and increased sensorimotor connectivity in the time window following practice (Deleglise et al., 2023). In line with these findings, we demonstrate greater ‘fast’ learning during initial learning of an implicit motor sequence is associated with both increased default mode network connectivity and decreased sensorimotor GABA concentration. In line with our analysis of micro-offline periods, this finding indicates that explicit sequence awareness may not be a necessary condition for involvement of hippocampal networks across early learning, aligning with models which propose that the hippocampus performs a more general ‘event sequence generating’ role in memory formation (Buzsáki & Tingley, 2018).

When examining learning outcomes across the entire practice period (i.e., 32 task blocks), we observed that the total degree of learning was linked with greater motor network connectivity. These findings add to a growing body of literature demonstrating interactions between hippocampal and motor networks in the initial ‘fast’ phase of motor learning (Buch et al., 2021; Deleglise et al., 2023; Jacobacci et al., 2020) that has traditionally been linked to interactions between cortico-striatal and cortico-cerebellar systems (Dayan & Cohen, 2011). Using a combination of MR spectroscopy and fMRI, we also provide evidence linking tonic GABA inhibition to motor network functional connectivity strength. Overall, our findings indicate implicit sequence learning as a function of local inhibitory and large-scale network connectivity strength. Future work is required to establish how these mechanisms may generalise to acquisition of more complex skills, and/or are implicated in consolidation and stabilisation processes across longer timeframes.

### Future directions and conclusion

In this study, we demonstrate that both default mode and motor network functional connectivity strength underlie the development of implicit motor skills. We show that the early phases of learning relate to local sensorimotor inhibition, which may be modulated via engagement in high-intensity interval exercise. However, there are certain methodological limitations which should be considered when interpreting these findings. First, we report correlational evidence linking network connectivity and GABA concentration to learning outcomes. Future work using dynamic causal modelling approaches or perturbation techniques (e.g., non-invasive brain stimulation methods) would be informative in establishing the causal contribution of particular regions (e.g., hippocampus) across temporal phases of implicit sequence learning. Second, we acquired resting-state fMRI at the end of the experimental protocol. We acknowledge that measuring resting-state connectivity at baseline would have eliminated any possible influence of the exercise protocol. However, to ensure we obtained valid MR spectroscopy estimates it was necessary to avoid any impact of gradient coil heating from the resting-state fMRI echo planar imaging sequence (Hui et al., 2021). We also did not observe any significant differences in resting-state connectivity between exercise groups, increasing our confidence that our results have not been inadvertently influenced by this aspect of the experimental design. Third, we focussed our analysis on the effect of exercise on motor learning across practice. However, our work and that of others suggests that exercise may have greater impact on the *retention* of motor skills relative to acquisition (Roig et al., 2012; Stavrinos & Coxon, 2017; Taylor et al., 2024). Future work may consider the addition of retention tests across longer timeframes to establish time-dependent effects of HIIT-priming on implicit learning.

Overall, via analysis of multiple neuroimaging modalities, we provide novel insights into the network and neurochemical mechanisms supporting implicit motor skill learning. High-intensity interval exercise is an effective method of modulating tonic concentrations of sensorimotor GABA, but may not lead to consistent facilitation of implicit sequence learning. Future research is required to elucidate causal relationships between network nodes and implicit learning across temporal scales.

